# More persistent bacterial than fungal associations in the microbiota of a pest

**DOI:** 10.1101/2022.02.28.482158

**Authors:** Kiran Gurung, Stefanie Nicoline Vink, Joana Falcão Salles, Bregje Wertheim

## Abstract

The invasive fly *Drosophila suzukii* is a pest that can infest a diverse range of intact, ripening fruits, using its serrated ovipositor. This constitutes a different niche compared to the rotting fruits its ancestors use, especially because these intact fruits have limited quantities of microbes and soluble nutrients for the developing larvae. To investigate the potential role of microbial associations in the niche expansion of this invasive fly, we characterized the bacterial and fungal communities of *D. suzukii* and various wild fruits from which they developed. To assess cross-generational microbial associations, we also lab-reared fly populations and characterized their microbial communities. Diversity metrics of microbial communities differed significantly between flies and fruits. Different fruit types varied substantially in microbial composition, while flies showed relatively uniform bacterial communities, irrespective of the fruit source they developed on. After lab-rearing, bacterial communities still showed considerable overlap with those of wild flies. Fungal communities of flies and fruits showed larger resemblance, with a substantial overlap between wild flies and the corresponding fruits on which they had developed. Our study thus reports that the fungal community structure in these pests largely reflects those on the breeding substrates, while these flies might have formed more persistent associations with bacteria and transmit these across generations rather than obtaining them from their food source.

## Introduction

The microbial communities associated with insects are highly diverse, and the dynamics within and between members of the microbiome can affect the fitness and behavior of insects in various ways (Gurung et al., 2019). Studies on insect microbiota have demonstrated that various factors shape the microbial community composition, ranging from life stages, environment, host genetics and diet (Yao et al., 2019). In some cases, microbes also contribute towards the pest status of invasive insects (Lu et al., 2016). When pest insects are (at least partially) dependent on microbes for their performance or fitness, these microbes have the potential of being utilized as pest management tools, and often one of the first steps is examining the microbiota profiles and their associations in the pests (Crotti et al., 2012; Lewis et al., 2019, Sacchetti et al., 2019, Yao et al., De Cock et al., 2020).

A pest of ripening fruits, *Drosophila suzukii* has its origin in Asia (*<u>Drosophila suzukii</u>*<u>, CABI. 2019</u>), and has successfully invaded several parts of America and Europe in the last decade (Calabria et al., 2012; Cini et al., 2014). The female lays eggs on intact ripening fruits and damages them, which is often accompanied by fruit tissue collapse, due to both feeding of the developing larvae inside the fruit, and the rot that is frequently induced by the infestation with *D. suzukii* (Ioriatti et al., 2018). Fruits become unsuitable for sale, resulting in huge economic losses for fruit growers (Goodhue et al., 2011; Atallah et al., 2014). *Drosophila suzukii* infests a large range of soft fruits, cherries, and different types of berries (Walsh et al., 2011; Rota-Stabelli et al., 2013). The ability of *D. suzukii* to colonize intact fruits is driven by its serrated ovipositor, which allows it to puncture the skin of fruits. This is a morphological innovation that enabled the species to expand from its ancestral niche of rotting and decomposing fruit to also infest fresh and ripening fruits (Atallah et al., 2014).

Ripening fruits have fewer nutrients (proteins and sugars) (Silva-Soares et al., 2017) and also harbor a more limited number of microbes than fermenting fruits. This poses possible dietary challenges to the larvae of *D. suzukii* that develop mostly inside the fruit host. A number of *Drosophila* species depend on yeasts during their larval stage for essential nutrients, and feed on these yeasts when they are developing on fruits (Starmer &Fogleman, 1986; Carvalho et al, 2010). In line with this, earlier research demonstrated that germ-free *D. suzukii* larvae failed to develop on protein-poor and fruit-based artificial diets, but this could be rescued with the supplementation of microbiota (Bing et al., 2018). If the microbial communities on intact fruits are indeed insufficient to sustain larval development, then during oviposition *D. suzukii* females may possibly inoculate the fruits with a set of microbes to supply their developing offspring with the required microbiota. Association with some microbes may thus have an essential impact on the life history of these flies. Furthermore, microbes vectored by the flies may have plant pathogenic potential, causing the fruit collapse (Hamby and Swett, 2015). The question as to what kind of microbial associations these pests might have can thus provide us with a better understanding of microbial contribution in the niche expansion of this invasive pest, in its broad life history, as well as in its pest status.

One of the earliest studies on the bacterial community composition in field-caught *D. suzukii* revealed *Tatumella* to be a dominant bacterium across larval and adult stages (Chandler et al., 2014). Although their exact role on insect host fitness is not yet known, the species *T. ptyseos* reportedly showed plant pathogenic traits (Marín□Cevada V et al., 2010). Furthermore, bacteria belonging to the families of *Acetobacteriaceae, Enterobacteriaceae* and *Firmicutes* have also been reported in *D. suzukii*. Some of these bacteria are associated with *Drosophila* in general and provide fitness benefits (Bing et al., 2018). Also yeasts such as *Hanseniasporum, Pichia*, and *Issatchenkia*, impact various life history traits of *D. suzukii* (Lewis and Hamby, 2019). Such reports on various microbial associations, point out a question whether the pest associates with microbes through exposure on their breeding site, or whether they might be retaining and transmitting microbiota in a more persistent association.

To investigate the microbial associations of this invasive fly, we characterized the similarities and differences among the bacterial and fungal communities of *D. suzukii* and various wild fruits from which they developed. The microbial communities that are typically associated with surfaces of various fruits and vegetables can vary considerably (Leff and Fierer, 2013). When developing larvae rely mostly on the microbial communities that they encounter on these fruits, we would expect that emerging flies from different fruits would exhibit high variations in their community patterns and align with those of the corresponding fruits. Alternatively, when *D. suzukii* formed more persistent associations with some microbes that they inoculate into the fruits while ovipositing, we would expect flies emerging from different fruit types to harbour similar microbiota. To explore whether emerged flies exhibit consistent microbial associations across various fruits, or whether fruit source plays an important role in determining the flies’ microbiome, we sampled the microbiome composition (bacteria and fungi) of fruits and the corresponding emerging *D. suzukii* flies, collected at four different locations in the Netherlands. Additionally, to identify members that potentially have a more intimate relationship with *D. suzukii*, and are vertically or horizontally transmitted, we subjected some of these wild flies to lab-rearing conditions and evaluated the changes in their microbiome after several generations.

## Methods

### Sampling wild flies and fruits

We collected fruits that were infested with *D. suzukii* in The Netherlands in 2018, across four different locations (between 7.5 - 200 km apart) and five fruit types (Summary in table 1). In three of these locations, we sampled multiple different fruit types or cultivars. Infestation was determined by identifying the dented spots on the fruits that resulted from egg laying. A large number of collected fruits were individually placed in plastic cups, parafilm-sealed and brought into the lab to further incubate them in bottles with autoclaved sawdust (saw dust soaks up unwanted moisture from decomposing fruits) until the adult flies emerged. Incubation conditions were 20 ^°^C, 65% relative humidity and 16 hr:8 hr light:dark cycle. The emerging flies (referred as “wild flies”) from each fruit and the corresponding fruits were collected and stored at -80^°^C until DNA extraction. An additional batch of flies that emerged from the same cherry, strawberry or elderberry fruits was subsequently lab-reared (referred as “lab-reared flies”).

**Table 1.** Details of the infested fruit collections (these fruits and the flies that emerged from them were used for microbiome characterization)

| Name | Location co- | Type of | Code by fruit | Fruit type | Whether |
| --- | --- | --- | --- | --- | --- |

|  | **ordinates** | **sampling**<br>**site** | **source** |  | **flies were**<br>**lab-reared** |
| --- | --- | --- | --- | --- | --- |
| Randwijk | N51°56.953'<br>E005°43.041' | Orchard | Cherry_Rw_1 | Cherry | Yes |
|  |  | Orchard | Cherry_Rw_2 | Cherry | No |
|  |  | Orchard | Cherry_Rw_3 | Cherry | No |
| Winssen | N51°53.234'<br>E005°41.372' | Orchard | Cherry_Wn_4, | Cherry | No |
|  |  | Orchard | Raspberry_Wn | Raspberry | No |
| Dirksland | N51°44.63'<br>E4°5.48' | Orchard | Raspberry_DI, | Raspberry | No |
|  |  | Orchard | Strawberry_DI, | Strawberry | Yes |
|  |  | Orchard | Blackberry_DI | Blackberry | No |
| Hoogeveen | N52°44.61'<br>E6°32.66 | Farm field | Elderberry_Hv | Elderberry | Yes |

<sup>1</sup> Each fly replicate corresponds to a pool of 5 adult flies that emerged from the same individual fruit; each fruit replicate corresponds to approximately 1g of fruit.
<sup>2</sup> Cherries from location Randwijk belonged to three different (and unknown) cultivars

### Lab rearing

We started three lab cultures from wild flies that emerged from 1) cherry collected in Randwijk, 2) strawberry collected in Dirksland, and 3) elderberry collected in Hoogeveen. These strains were reared in the lab on artificial food (see supplementary file for the fly food composition), for approximately 10 generations, before being collected and stored at -80□C for microbiome characterization. The fly food contained sucrose, glucose, agar, cornmeal, wheatgerm, soy flour, molasses, yeast, ethanol, and was supplemented with antimicrobial agents (propionic acid and nipagin) to reduce mold contamination. The founding population of flies was composed of 8-10 individuals per fruit type. Each successive generation, at least fifty individual flies were transferred to the fresh medium.

### DNA extraction and sequencing

We collected five biological replicate samples for each fruit type: Each fly sample consisted of five female flies (wild and lab-reared flies) that emerged from a single fruit, and approximately 1g of fruit was excised with sterilized bladed for the corresponding fruit samples. DNA extractions were performed using Qiagen Soil DNA extraction kit according to manufacturer’s protocol. Prior to extraction, pooled flies in 2mL centrifugation tubes were washed once in sterile water by vortexing for approximately 10 seconds to remove saw dust. Finally, in order to check for any kit-associated contamination (Glassing et al., 2016), five extraction controls with no samples were subjected to DNA extraction and further sequencing.

Following extractions, we quantified DNA using NanoDrop2000 (Thermo Fisher Scientific, MA, USA). Sequencing was outsourced to the University of Minnesota Genomics Center (USA), on Illumina MiSeq platform (V3 Chemistry and 2×300 paired end run), targeting the V4-V6 region of the 16S rRNA gene and ITS1 region for bacteria and fungi respectively (Primer sequence information in supplementary file).

### Sequence analyses

We processed the sequences in QIIME2 platform (version 2019.10, Boylen et al., 2019). To avoid losing a substantial amount of reads after pairing (due to non-overlapping coverage of the amplicon), we chose to analyze only the forward reads for both bacteria and fungi. For bacteria, we trimmed the primers using the ‘Cutadapt’ plugin (Martin, 2011). We denoised and chimera filtered the sequences using DADA2 plugin with its default settings at phred score of at least 25 and truncation length of 220 bp (Callahan et al., 2016). Taxonomic assignment was done using RDP classifier (Wang et al., 2007). We removed sequences belonging to archaea, eukaryotes, chloroplast and mitochondria. For fungi, we used ITSxpress plugin (version 1.7.2, Rivers et al., 2018) to extract the fungal ITS1 region. Denoising and chimera filtering was performed using DADA2 with a truncation length of zero. Taxonomic assignment was performed using the UNITE database (version 8_99_04.02.2020, Abarenkov et al., 2010) that was trained and classified. We used the feature table and taxonomy table for both bacteria and fungi and additionally phylogenetic tree for bacteria generated from the QIIME2 workflow in R (version 3.6.3, R Core Team). The reads in the extraction controls were subtracted and removed from the read abundances in the samples prior to downstream analysis (taxa found in controls are provided in the supplementary file).

### Statistical analysis

After excluding controls, there were a total of 105 samples that comprised fruits, wild flies and lab-reared flies. This whole dataset was used to provide an overview of the microbial communities associated with flies and fruits. Comparative analyses were done on subsets: for the fruit-wild fly comparisons (90 samples, bacterial and fungal community) and for the lab-reared fly-wild fly comparisons (30 samples, bacterial community). We did not include the analyses from fungal communities of lab-reared and wild fly subsets as we used antifungal agents which could be a highly confounding factor in their community pattern, but we have added the data in the supplementary file.

For the fruit-wild fly comparisons, the data were rarefied to a sequencing depth of 4500 for bacteria and 7800 for fungi, resulting in the loss of seven and six samples for bacterial and fungal communities respectively. For the lab-reared fly-wild fly comparison, the reads were rarefied to a sequencing depth of 4500 for bacteria with no loss of samples. Analysis on alpha and beta diversity were performed on rarefied data, while remaining analyses were performed on unrarefied data.

For bacterial alpha diversity we calculated richness (observed ASVs) and Faith’s phylogenetic diversity using packages phyloseq (version 1.30.0, McMurdie and Holmes, 2013) and picante (version 1.8.1, Kembel et al., 2010). Fungal alpha diversity was calculated based on richness and Shannon index using phyloseq. We used the Kruskal-Wallis test (Kruskal and Wallis, 1952) for comparing the alpha diversity between different sample types (i.e., the different types of fruit sources, and the different groups of emerging wild flies), followed by post-hoc Dunn tests with Benjamini-Hochberg p-adjustment method (Dunn, 1961; Benjamini and Hochberg, 1995) using phyloseq and dunn.test (version 1.3.5).

We determined the beta diversity by PCoA based on UniFrac distance metrics for bacterial communities (Lozupone and Knight, 2005) and Bray Curtis metrics for fungal communities (Beals, 1984) using the packages phyloseq and vegan (2.5-6, Oksanen, 2015). We tested for statistically significant differences among all the fruits-wild flies, and between lab-reared flies-wild flies using PERMANOVA (Anderson, 2014). Differences in community dispersion was checked by PERMDISP (Anderson, 2006) for the significant p-values resulting from PERMANOVA.

Using procrustes analysis, we further tested the degree of association of the bacterial and fungal communities based on the Bray Curtis metrics, between the fruits-wild fly subset as well as between the wild and lab-reared fly subset (Jackson, 1995), with codes adapted from van Veelen et al. (2020).

We assessed taxonomic relative abundances across fruits-wild flies and in the lab-reared flies-wild flies using package microbiome (version 1.12.0, Lahti and Shetty, 2017), with default parameters and transformation set to ‘compositional’.

## Results

### Microbial diversity of fruits and associated flies

The alpha diversity of bacterial communities across the fruits and wild flies differed by fruit types in terms of observed ASVs (*X*^*2*^=25.829, p-value < 0.001, Figure 1a) and Faith’s phylogenetic diversity (*X*^*2*^=12.085, p-value < 0.001, Fig.S1a). In general, alpha diversity was higher in wild flies than in fruits, except for the flies originating from raspberries in both locations Winssen and Dirksland (dunn test, p-value=0.05).

**Figure 1.**
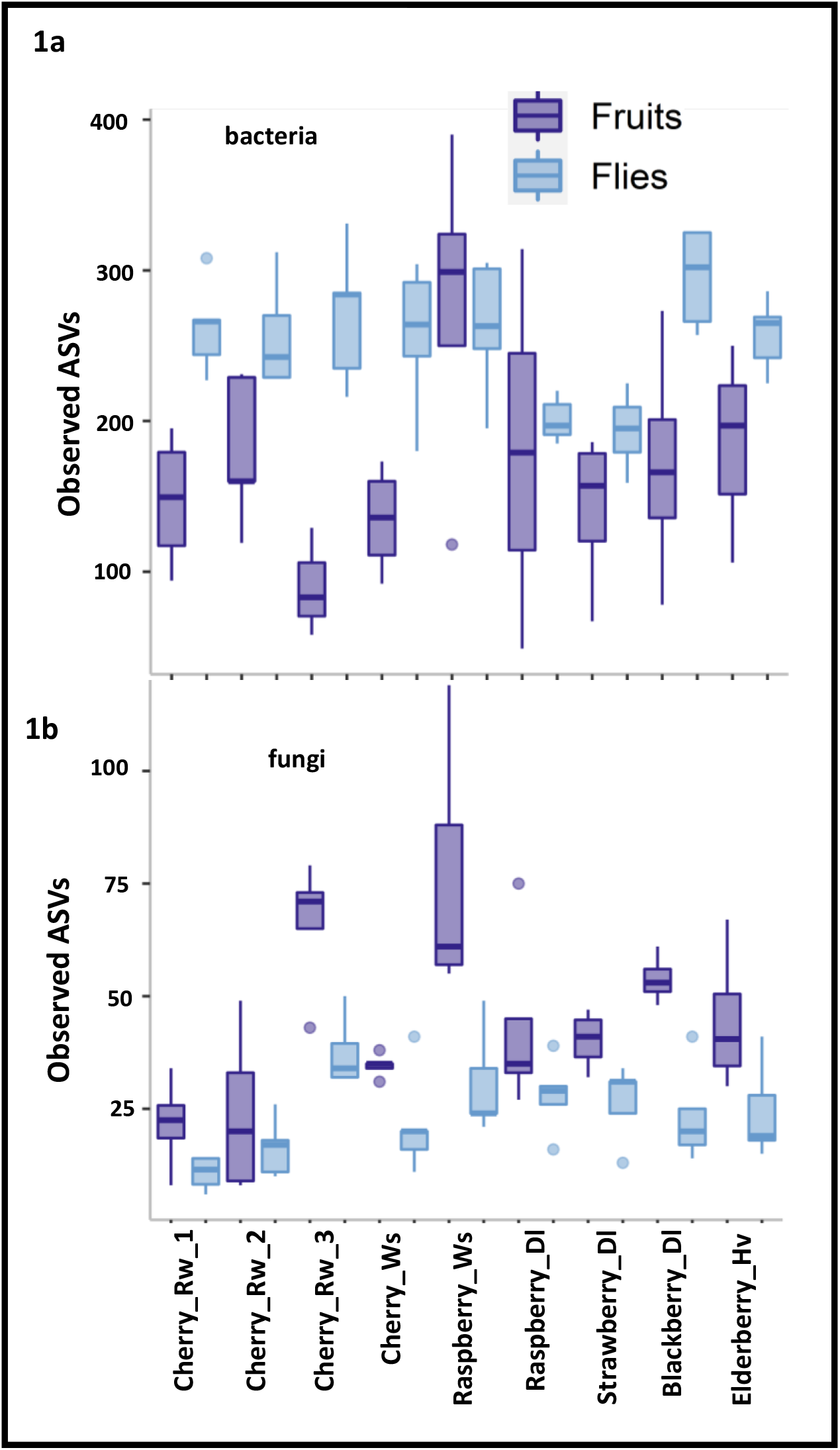
Alpha diversity (number of observed ASVs) of bacterial (a) and fungal (b) communities associated with fruits and wild *D. suzukii* flies, arranged by fruit type and collection site. The richness of both bacterial and fungal communities across fruits and the emerging wild flies differed significantly (p-values<0.1 and <0.05, respectively).

Richness and Shannon diversity index of fungal communities also differed significantly in fruits and wild flies according to the fruit types (*X*^*2*^=22.702, p-value <0.001, Fig 1b; *X*^*2*^=48.429, p-value < 0.001, Fig. S1b; dunn test, p-value=0.05), but contrary to richness for bacterial communities, the fungal communities on fruits generally showed higher alpha diversity than the flies.

### Microbial community patterns in fruits and associated flies

To assess whether the microbiota of the flies would reflect those of the fruits, we compared the microbiome composition of the fruit sources and wild flies (Figure 2). The bacterial communities of wild flies clustered together and away from their corresponding fruit sources, thereby indicating a strong separation across these two sample types. For bacterial communities, PERMANOVA based on weighted UniFrac distance revealed significant differences between the wild fly and their corresponding fruits (R^2^=0.2688, p-value=0.001, figure 2a &2b). Additionally, the communities were homogeneous in their dispersion (PERMDISP, p-value=0.755). Based on unweighted UniFrac, PERMANOVA (R^2^=0.115, p-value = 0.001, figure S2) showed significant difference and dispersion was not homogenous (PERMDISP, p-value = 0.001, figure S2). While replicate samples for the same fruit type per location clustered mostly together, they showed some variance both within and among fruit types (figure S2).

**Figure 2.**
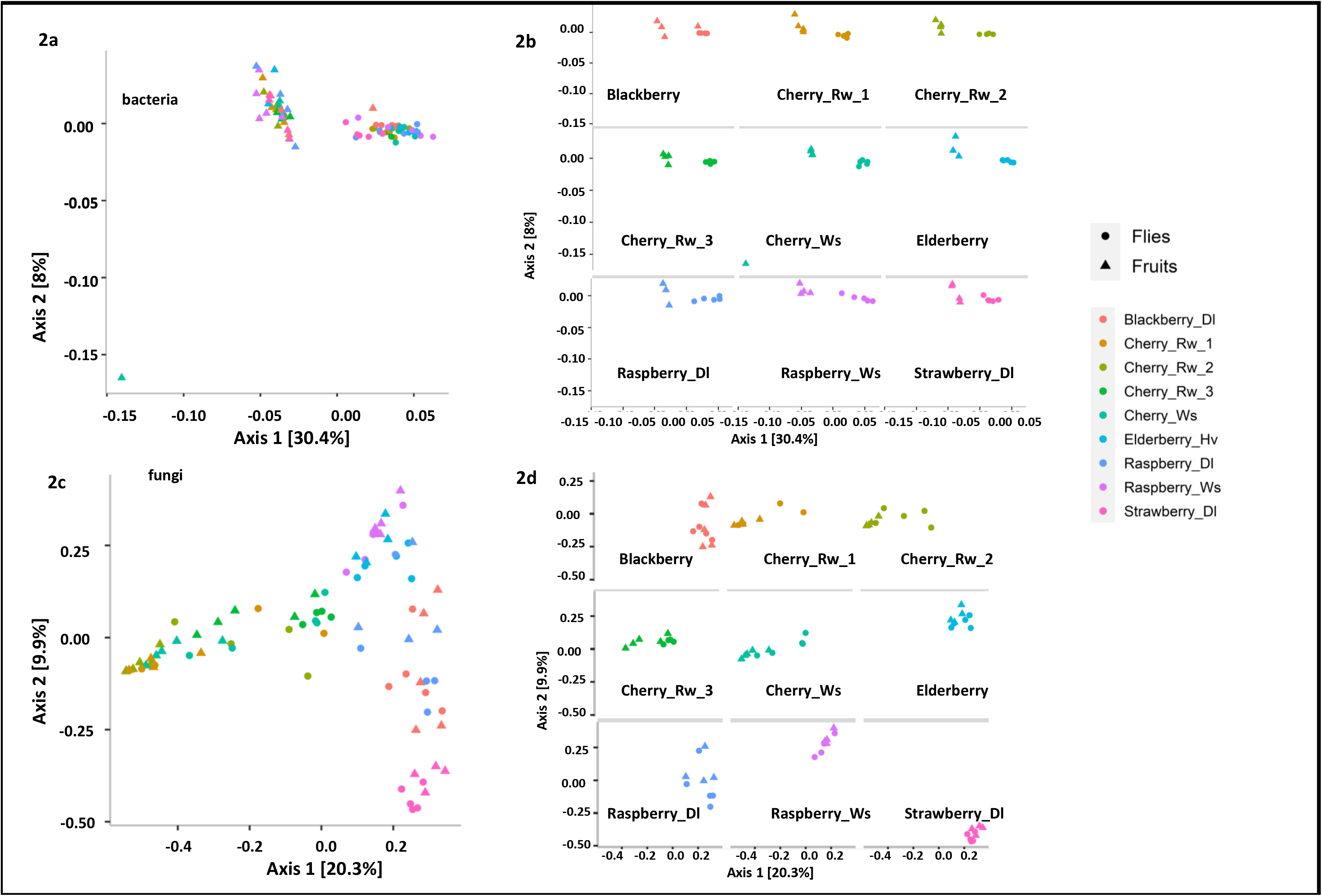
Beta diversity of bacterial (a and b) and fungal (c and d) communities associated with fruits (triangles) and wild *D. suzukii* (dots). **a:** PCoA of weighted UniFrac distance of bacterial communities differed significantly between fruits and flies (p-value=0.001). **b:** Individual facets of PCoA using weighted UniFrac for every fruit type in every collection site. **c:** PCoA of Bray Curtis dissimilarity index for fungal communities, which differed significantly between fruits and flies (p-value=0.021). **d:** Individual facets of PCoA using Bray Curtis by fruit type in every collection site.

The beta diversity of fungal communities, assessed using PERMANOVA on Bray-Curtis distance, also showed differences between wild flies and their corresponding fruits (R^2^=0.022, p-value=0.021, figure 2c &2d). Unlike the bacterial communities, however, the fungal communities showed no clear clustering into either fruit or fly samples (Figure 2d); rather the fruit and fly samples overlapped considerably within fruit type, but fruit types differed substantially from each other. The dispersion was heterogeneous (PERMDISP, F-value=8.30, p-value=0.005), revealing differences in within-group variances. We observed considerable overlap of the fungal communities between the wild fly and its corresponding fruit source, as shown from the clustering by fruit source in most of the combinations (Figure 2d). Additionally, considerable clustering occurred between different fruits and flies from similar locations (as noted for the three cherry cultivars from Randwijk, and among strawberries, raspberries and blackberries from Dirksland).

We further performed procrustes analysis to assess if the microbial communities across sample subsets covaried (Table 2). For the bacterial communities, we did not observe concordance between the wild flies and the corresponding fruit subset (M^2^=0.934, p-value=0.19). In contrast, association between fungal communities of wild flies and corresponding fruits was significant (M^2^ = 0.554, p-value=0.001).

**Table 2.**
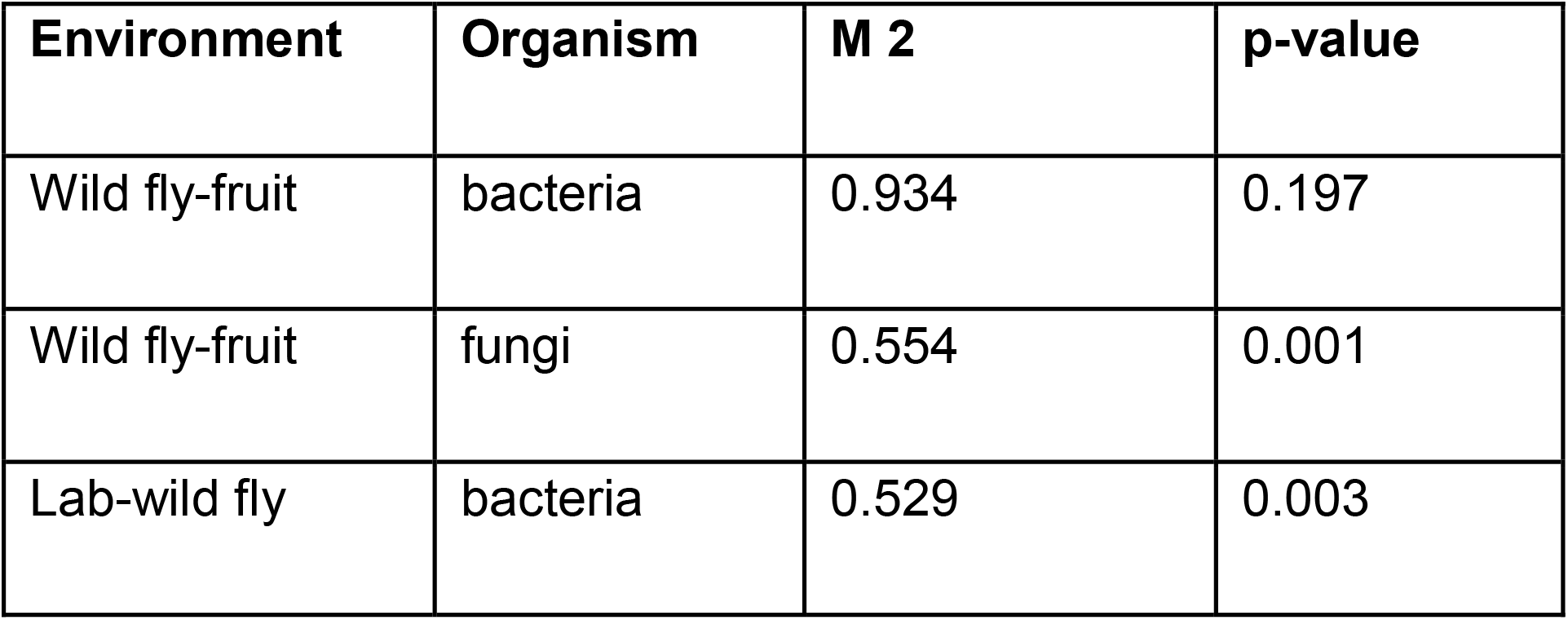
Procrustes analysis indicating the association between the bacterial and fungal communities from wild flies and their corresponding fruit samples as well as between lab flies and wild flies.

### Bacterial community across wild and lab-reared flies

Bacterial richness in lab-reared flies was similar to that of their wild counterparts, except for the samples from strawberry, where lab-reared flies showed a higher alpha diversity (figure 3a). Overall, we did not find any difference between the wild and the lab flies (p-value=0.40). Also, the Faith’s phylogenetic diversity indices did not differ significantly across the sample types (p-value=0.101, figure S3).

**Figure 3.**
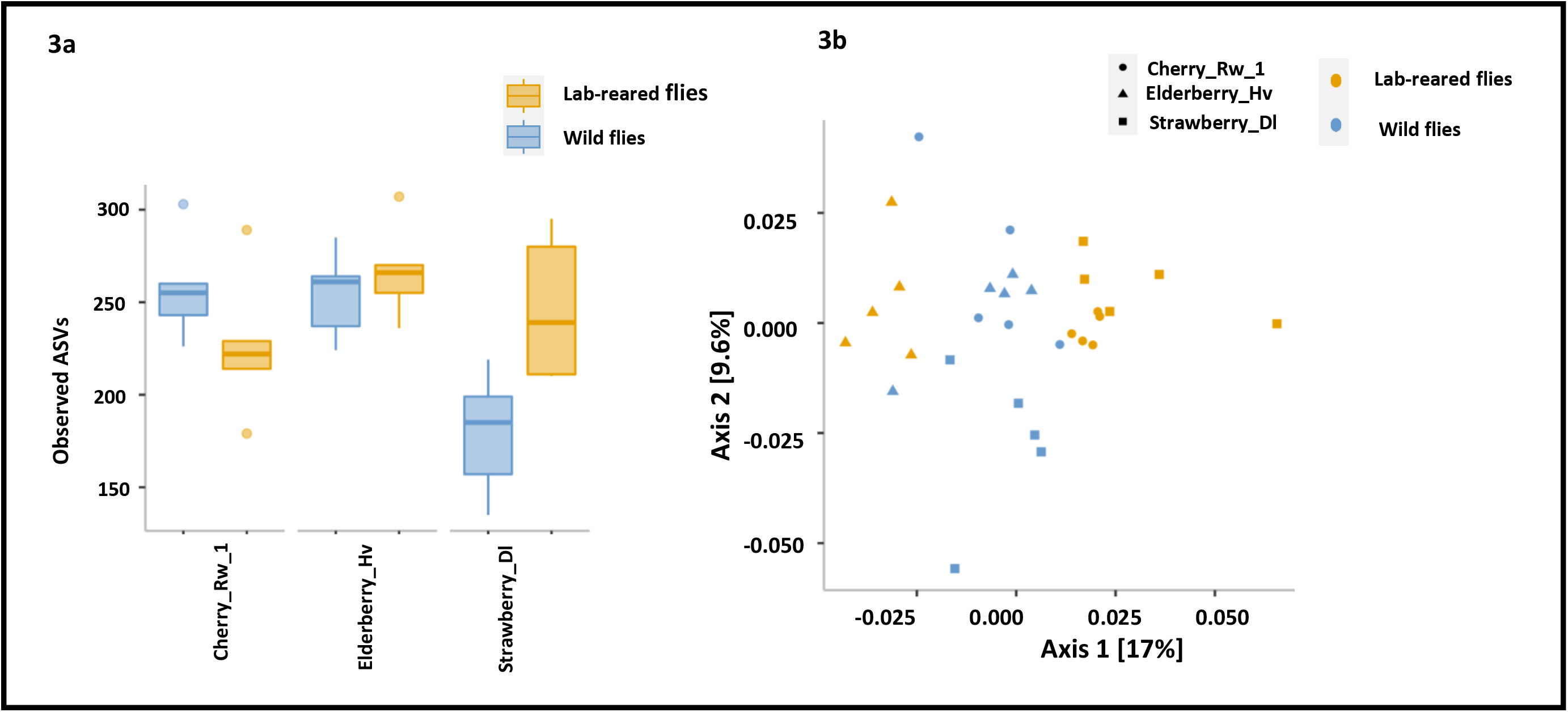
Alpha and beta diversity (of bacterial communities across wild and lab-reared flies. Alpha diversity (observed ASVs) of a) did not differ significantly between wild and lab-reared flies (p-value=0.40). Beta diversity c) based on Weighted UniFrac distances for bacterial communities differs significantly between the lab-reared and wild flies (p-value=0.001)

We observed significant differences in beta diversity of the bacterial communities between lab-reared flies and their respective wild flies in composition (R^2^=0.06, p-value=0.011, figure 3b), but only 6% of the variation could be explained by the rearing environment. We did not observe any differences in the dispersion (PERMDISP, F-value=3.15, p-value=0.66). Further, the correspondence in bacterial communities between wild flies and lab-reared flies was significant (M^2^ = 0.5298, p-value=0.003, Table 2).

### Community composition and core microbes among different sample types

We profiled the relative abundances of top-10 most abundant bacteria and fungi across the different samples (Figure 4). At the genera level, we observed *Candidatus Carsonella, Ca. Scalindua, Gluconobacter, Minicystis, Omnitrophica genera incertae sedis, Pantoea, Pelospora, Stella, Tatumella, Tephidisphaera* among the top 10 abundant bacterial genera across the wild flies and fruit samples. Some of these genera were limited to only the flies, or were recorded at a higher abundance in flies than in the corresponding fruits (Figure 4a). We further identified four bacteria that were found in at least 95% of wild fly samples at an abundance threshold of 0.001 (Figure 4b). These were *Succinatimonas, Pelospora, Ca. Carsonella* and *Ca. Omnitrophica*. In the fruit samples, we were unable to identify any bacterial genera that were common to all the fruit samples, even at lower prevalence. Furthermore, these fly core bacteria were only retrieved from a few fruit samples and in much lower abundances than in wild flies, except for one fruit sample (figure S4). Three of these core bacteria were also observed in the lab-reared flies (Figure 4C). In addition, we found *Acetobacter* in the lab-reared flies, which was missing in most of the ancestral wild flies emerging from cherries and elderberries.

**Figure 4.**
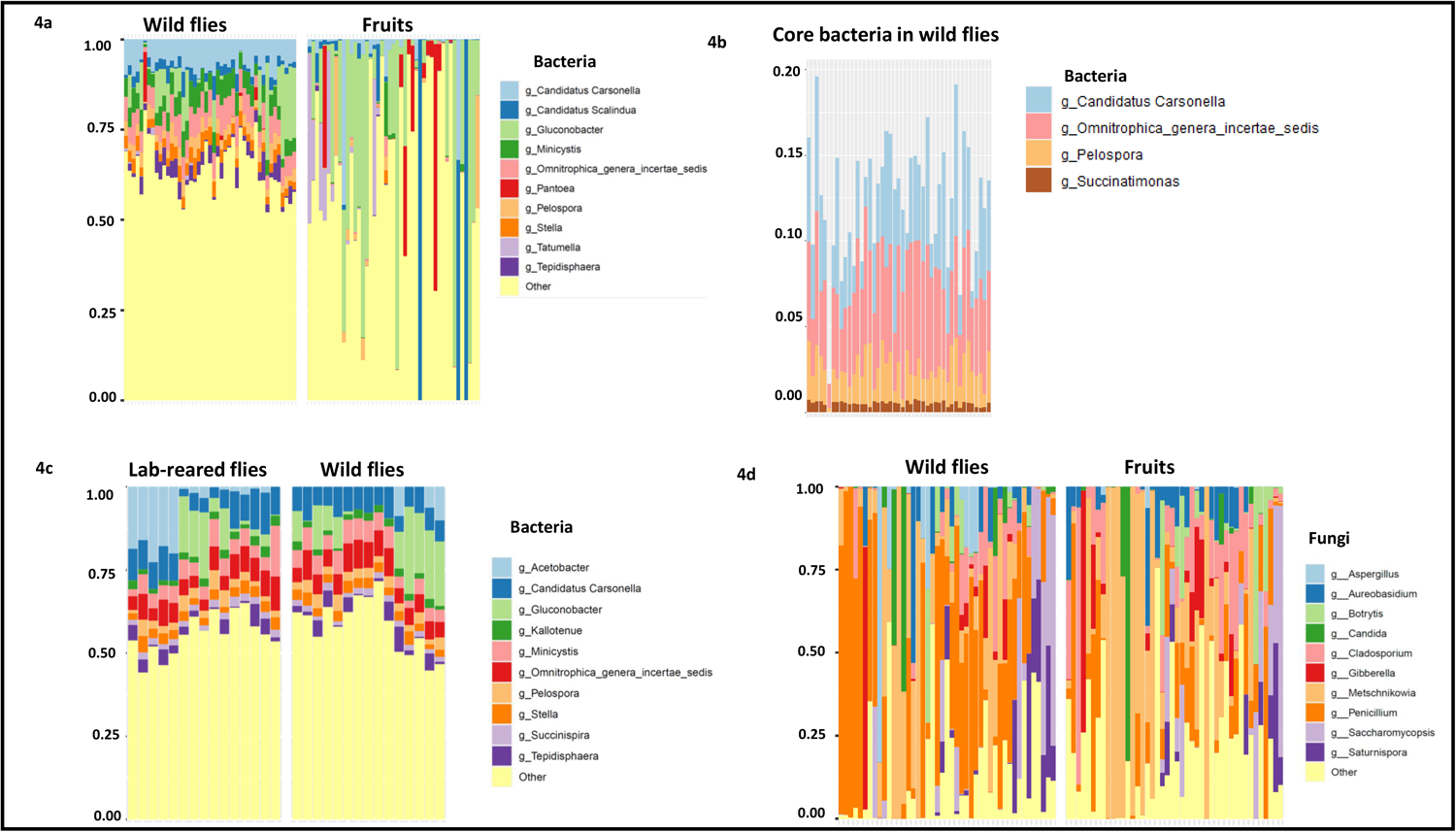
Top 10 abundant bacteria across wild flies and their source fruits (a). Four core bacteria were observed across the wild flies (b). Top 10 abundant bacteria across lab-reared flies and wild flies, which also comprised three of the core bacteria (c). Top 10 abundant fungi across wild flies and their corresponding fruits (d).

Both yeasts and molds made up the fungal composition across the fruit and wild fly subsets, mostly belonging to the genera *Aureobasidium, Saturnispora, Botrytis, Candida, Cladosporium, Saccharomycopsis, Gibberella, Metschnikowia, Penicillium*, and *Aspergillus*, with the latter three also being present in the lab-reared flies (Figure 4d and S5c). We did not find any core fungal ASVs at 95% prevalence across either fruits or wild flies, or across both combined. Furthermore, there was no core fungal ASV common in the wild fly subset even with relaxed criteria of lower prevalence and detection threshold. In the fruit samples, however, we did find *Mycosphaerella tassiana* at a prevalence of 70% and 0.0001 abundance threshold. Hence, almost all wild flies shared a subset of bacteria, and occurrence of some of these bacteria was rare in fruits. In contrast, fungi were not commonly shared among the wild flies.

## Discussion

In this study we characterized the bacterial and fungal communities associated with wild *D. suzukii* flies, an invasive pest insect originating from Asia, which causes severe damage in the production of soft fruits around the globe (Walsh et al., 2011). The success of this species lies in its ability to perforate the fruit skin to lay eggs on ripening fruits, as well as its capacity to develop in a large variety of these fruits. This new niche of ripening fruits may lack essential nutrients that microbes could be providing (Bing et al., 2018). Our primary focus was to study the microbiome in *D. suzukii*, as well as its associated fruit substrates, to assess whether the flies acquired their associated microbiota mostly from exposure to the microbial communities of the fruits on which they developed - in this case, we expected substantial overlap of the fruit and fly microbiota with differences in microbial composition for flies emerging from different fruit types - or whether they have formed more persistent associations with some microbiota that they retain irrespective of their fruit hosts.

We observed significant differences in bacterial and fungal diversity among different fruit types, but the flies that emerged from these fruits had a remarkably uniform bacterial community composition. In contrast, the fungal community distribution in the flies largely resembled those in their host fruits. Flies that were lab-reared for 10 generations still harboured some of the bacterial genera that largely resembled those of their wild ancestors. We observed no significant difference in alpha diversity metrics between bacterial communities of lab-reared and wild flies. Perhaps this could also indicate that lab-rearing for 10 generations might not have been sufficient to change the community pattern. For fungal communities, although we observed no significant difference in beta diversity between the lab-reared and wild flies, we did see a severe reduction in alpha diversity for lab-reared flies, but this is likely attributable to the use of anti-fungal agents (i.e., proprionic acid and nipagin) in the artificial diet (Figure S5a and S5b). Thus, the bacterial communities in emerging flies were largely shaped irrespective of the fruit type and are distinct from the bacterial communities on the fruits itself. In contrast, the fruit source most likely potentiated the fungal community distribution in the flies. At the level of ASVs, several of the microbes present in the wild flies also seem to be present in the lab-reared samples, suggesting that they were vertically or horizontally transmitted.

It is important to notice that in our study we did not analyze the larval microbiome, as we were interested specifically in the associated microbes of the adults. In holometabolous insects like *Drosophila*, metamorphosis induces changes at morphological and physiological level, which restructures several tissues and might also result in shedding of microbes (Tissot and Stocker, 2000; Johnston and Rolff, 2015). However, sometimes, depending on the family or life history of the insects, a few selected microbes might be retained (Johnston and Rolff, 2015). In order to verify whether such restructuring during development results in microbial shifts for *D. suzukii*, future studies should also focus on determining the microbiome of wild *D. suzukii* larvae as well as pupae collected from fruits across a broader geographical range. In line with our findings, another study on associated microbes of the *Bactrocera* tephritid fruit flies showed difference in bacterial community pattern for fruits and larvae (Majumder et al., 2019). In *Drosophila* species, Martinson et al. (2017) reported the bacterial community composition in flies to be distinct from their source food. Although we do not yet know how this separation of fruit and fly bacterial communities can persist, the continuation of the bacterial associations for 10 generations on artificial diet suggest that perhaps ovipositing females can transmit bacteria across generations to their offspring. And in addition to diet, host type may also have been a factor driving the community composition in our fly samples.

*Drosophila suzukii* flies spend a substantial amount of time in close association with the fruits from which they emerge as adults. We therefore expected to observe a subset of microbes commonly shared across both fruits and flies. At the genera level, we observed *Gluconobacter, Pantoea, Tatumella* across the fruits and some of the wild flies, as reported by previous studies (Chandler et al., 2014; Vacchini et al., 2017; Solomon et al., 2019). We also observed bacteria *Ca. Carsonella, Ca. Scalindua, Succinatimonas etc*. to be more abundant in the flies, and *Acetobacter* to be also present in the lab flies. Some of the prevalent fungal members of the genera *Penicillium* and *Metschnikowia* were common among wild flies and fruits, with the former being dominant in the lab-reared flies as well. In addition, few other fungi like *Pichia, Botrytis, Cladosporium* were also noted in some of the fly samples. However, no fungal genera were consistently shared among wild flies or fruits. The fungal community composition varied among fruit types, and those of the emerged flies showed substantial overlap. Similar to our findings, previous studies on coleopteran insects also suggested a strong role of the immediate environment in structuring the fungal communities in the insects (Kudo et al., 2019; Rassati et al., 2019).

Understanding the interactions between microbiome and insects is interesting from an evolutionary perspective, as it may facilitate niche expansions and adaptations to challenging environmental conditions. Furthermore, the core bacteria that we observed in our samples reportedly engage in carbohydrate and amino acid metabolic pathways (Riley et al., 2017; Morotomi et al., 2010; Stackebrandt and Hespell, 2006; Matthies et al., 2000). In the case of pest insects, it is also highly relevant to understand what determines the composition of these microbial communities, as this may reveal unknown aspects on their fundamental biology, sometimes also providing insight into vulnerabilities that can be exploited in pest management strategies (Raza et al., 2020). Culture based and sequencing approaches have revealed the influence of environmental factors on the bacterial community composition of pest insects, such as diet, climate and geographic location (Rizzi et al., 2013; Jones et al., 2019; Colman et al., 2012). To understand the processes by which associations between pest insects and microbiota arise, it is important to investigate the microbial composition on their food source (Behar and Jurkevitch, 2008; Beaulieu et al., 2017), which has been reported as a potential source from which insects can acquire their associated microbiota (Jones et al., 2019). Yet, it is also important to realize that specific insect-microbe associations can arise that transcend the intermediate of a food source.

The ability of *D. suzukii* to grow on such a wide range of fruits makes it an interesting system to explore whether fruit source plays an important role in determining its microbiome. Microbial communities vary considerably among various fruits and vegetables (Leff and Fierer, 2013), as we corroborated in the analyses of microbial communities on our fruit samples. We speculated that flies developing on different fruits would exhibit high variations in their community patterns and align with those of the corresponding fruits when they relied on their fruit hosts for acquiring microbes. Our results, however, showed that the bacterial communities of the wild flies were largely distinct from those of the fruits, and showed large similarities irrespective of fruit type.

Some of the microbial members that were found in wild flies from different fruits were also retained after 10 generations of lab-rearing. Based on these findings, we propose that *D. suzukii* may have formed persistent associations with some of these bacteria. In order to investigate what precisely fosters the host-microbiome interactions in this pest, studies similar to Ben-Yosef et al. (2014) could likely answer whether indeed there is any particular microbial associations that drives the pest status of *D. suzukii*. The fungal communities seemed to be less persistent, and here we did see more alignment in community patterns between fruits and emerging flies. This does not necessarily imply that these fungi are unimportant in the niche expansion, as they could be providing essential nutrients, e.g., sterols (Carvalho et al, 2010). However, it does appear that the associations with fungi are more transient than those with bacteria.

Concluding, our study reports the microbial community structure of *Drosophila suzukii* across a range of fruits. By characterizing the microbial communities of both the flies and the fruits from which they emerged, we could assess the similarity and differences between these communities across the fruits and the flies, in several different fruit sources. The characterization of these microbial communities represented, thus, an important step to identify any potential symbiotic relationship between these flies and bacterial or fungal species

## Acknowledgments

We thank Herman Helsen who provided the details of infestation and the growers. We are grateful to Tom Groot (Dirksland), Maria Buitenkamp and Edo Biewinga (Hoogeveen); Gijs Gerritse (Randwijk); Jeroen Spitzen (Winssen) who provided access to their garden and orchards. This work was supported by the Adaptive Life scholarship program (2017) awarded to KG by the University of Groningen. We thank the Center for Information Technology of the University of Groningen for their support and for providing access to the Peregrine high performance computing cluster. This research has been carried out in the groups of Evolutionary Genetics and Microbial Ecology at the Groningen Institute for Evolutionary Life Sciences (GELIFES) according to the requirements of the Graduate School of Science and Engineering (Faculty of Science and Engineering, University of Groningen; Groningen, the Netherlands).

## Author contribution

KG, JFS, BW conceived the experimental plan. KG performed all analyses. SNV and JFS provided support with sequence analysis. BW provided support with statistical analysis. KG wrote the original draft. KG, SNV, JFS and BW reviewed and edited subsequent drafts.

## Data availability statement

Additional files and data available in the link : https://figshare.com/s/4eebffffaab656b540f8

## References

Abarenkov, Kessy, et al. The UNITE database for molecular identification of fungi – recent updates and future perspectives. New Phytol 2010; 186.2: 281–285.

Anderson MJ. Distance-based tests for homogeneity of multivariate dispersions. Biometrics. 2006; 62(1):245–53.

Anderson MJ. Permutational multivariate analysis of variance (PERMANOVA). Wiley statsref: statistics reference online. 2014; 14:1–5.

Atallah J, Teixeira L, Salazar R, Zaragoza G, Kopp A. The making of a pest: the evolution of a fruit-penetrating ovipositor in Drosophila suzukii and related species. Proc R Soc Lond [Biol]. 2014; 281(1781):20132840.

Bauer DF. Constructing confidence sets using rank statistics. J Am Stat Assoc. 1972; 67(339):687–90.

Beals EW. Bray-Curtis ordination: an effective strategy for analysis of multivariate ecological data. Adv Ecol Res. 1984; 14:1–55.

Beaulieu M, Franke K, Fischer K. Feeding on ripening and over-ripening fruit: interactions between sugar, ethanol and polyphenol contents in a tropical butterfly. J Exp Biol. 2017; 220(17):3127–34.

Behar A, Jurkevitch E, Yuval B. Bringing back the fruit into fruit fly–bacteria interactions. Mol Ecol. 2008; 17(5):1375–86.

Benjamini Y, Hochberg Y. Controlling the false discovery rate: a practical and powerful approach to multiple testing. J R Stat Soc Series B Stat Methodol. 1995; 57(1):289–300.

Ben-Yosef M, Pasternak Z, Jurkevitch E, Yuval B. Symbiotic bacteria enable olive fly larvae to overcome host defences. R. Soc. Open Sci. 2015; 2(7):150170.

Bing X, Gerlach J, Loeb G, Buchon N. Nutrient-dependent impact of microbes on Drosophila suzukii development. MBio. 2018; 9(2).

Bolyen E, Rideout JR, Dillon MR, Bokulich NA, Abnet CC, Al-Ghalith GA, Alexander H, Alm EJ, Arumugam M, Asnicar F, Bai Y. Reproducible, interactive, scalable and extensible microbiome data science using QIIME 2. Nat biotechnol. 2019; 37(8):852–7.

Calabria G, Máca J, Bächli G, Serra L, Pascual M. First records of the potential pest species Drosophila suzukii (Diptera: Drosophilidae) in Europe. J Appl Entomol. 2012;136(1-2):139–47.

Callahan BJ, McMurdie PJ, Rosen MJ, Han AW, Johnson AJ, Holmes SP. DADA2: high-resolution sample inference from Illumina amplicon data. Nat Methods. 2016; 13(7):581–3.

Carvalho M, Schwudke D, Sampaio JL, Palm W, Riezman I, Dey G, Gupta GD, Mayor S, Riezman H, Shevchenko A, Kurzchalia TV. Survival strategies of a sterol auxotroph. Development. 2010;137(21):3675–85.

Chandler JA, James PM, Jospin G, Lang JM. The bacterial communities of Drosophila suzukii collected from undamaged cherries. PeerJ. 2014; 2:e474.

Cini A, Anfora G, Escudero-Colomar LA, Grassi A, Santosuosso U, Seljak G, Papini A. Tracking the invasion of the alien fruit pest Drosophila suzukii in Europe. J Pest Sci. 2014; 87(4):559–66.

Colman DR, Toolson EC, Takacs-Vesbach CD. Do diet and taxonomy influence insect gut bacterial communities? Mol Ecol. 2012; 21(20):5124–37.

Crotti, E., Balloi, A., Hamdi, C., Sansonno, L., Marzorati, M., Gonella, E., Favia, G., Cherif, A., Bandi, C., Alma, A. and Daffonchio, D. (2012). Microbial symbionts: a resource for the management of insect-related problems. Microb Biotechnol., 5(3), 307–317.

Currie CR. A community of ants, fungi, and bacteria: a multilateral approach to studying symbiosis. Annu Rev Microbiol. 2001; 55(1):357–80.

De Cock, M., Virgilio, M., Vandamme, P., Bourtzis, K., De Meyer, M., & Willems, A. (2020). Comparative microbiomics of tephritid frugivorous pests (Diptera: Tephritidae) from the field: A tale of high variability across and within species. Front Microbiol, 1890.

Dunn OJ. Multiple comparisons among means. J Am Stat Assoc. 1961; 56(293):52–64.

Folch PL, Bisschops MM, Weusthuis RA. Metabolic energy conservation for fermentative product formation. Microb Biotechnol. 2020.

Glassing A, Dowd SE, Galandiuk S, Davis B, Chiodini RJ. Inherent bacterial DNA contamination of extraction and sequencing reagents may affect interpretation of microbiota in low bacterial biomass samples. Gut pathog. 2016; 8(1):1–2.

Goodhue RE, Bolda M, Farnsworth D, Williams JC, Zalom FG. Spotted wing drosophila infestation of California strawberries and raspberries: economic analysis of potential revenue losses and control costs. Pest Manag Sci. 2011; 67(11):1396–402.

Gurung K, Wertheim B, Falcao Salles J. The microbiome of pest insects: it is not just bacteria. Entomol Exp Appl. 2019; 167(3):156–70.

Hamby KA, Swett CL. Elucidating symbioses between Drosophila suzukii and fungal communities for improved insect and disease management in raspberry production. North American Bramble Growers Research Foundation Funding. 2015.

Ioriatti C, Guzzon R, Anfora G, Ghidoni F, Mazzoni V, Villegas TR, Dalton DT, Walton VM. Drosophila suzukii (Diptera: Drosophilidae) contributes to the development of sour rot in grape. J Econ Entomol. 2018; 111(1):283–92.

Jackson DA. PROTEST: a PROcrustean randomization TEST of community environment concordance. Ecoscience. 1995; 2(3):297–303.

Johnston PR, Rolff J. Host and symbiont jointly control gut microbiota during complete metamorphosis. PLoS Pathog. 2015; 11(11):e1005246.

Jones AG, Mason CJ, Felton GW, Hoover K. Host plant and population source drive diversity of microbial gut communities in two polyphagous insects. Sci Rep. 2019; 9(1):1–1.

Kembel SW, Cowan PD, Helmus MR, Cornwell WK, Morlon H, Ackerly DD, Blomberg SP, Webb CO. Picante: R tools for integrating phylogenies and ecology. Bioinformatics. 2010; 26(11):1463–4.

Kruskal WH, Wallis WA. Use of ranks in one-criterion variance analysis. J Am Stat Assoc.1952; 47(260):583–621.

Kucuk RA. Bacterial Diversity of the Gut of Cotinis nitida. (2019).

Kudo R, Masuya H, Endoh R, Kikuchi T, Ikeda H. Gut bacterial and fungal communities in ground-dwelling beetles are associated with host food habit and habitat. ISME J. 2019; 13(3):676–85.

Lahti L, Shetty S. Tools for microbiome analysis in R. Version 1.1. 10013. 2017

Leff JW, Fierer N. Bacterial communities associated with the surfaces of fresh fruits and vegetables. PloS One. 2013; 8(3):e59310.

Lewis MT, Hamby KA. Differential impacts of yeasts on feeding behavior and development in larval Drosophila suzukii (Diptera: Drosophilidae). Sci Rep. 2019; 9(1):1–2.

Lu, M., Hulcr, J., & Sun, J. (2016). The role of symbiotic microbes in insect invasions. Annual Review of Ecology, Evolution, and Systematics, 47, 487–505.

Lozupone C, Knight R. UniFrac: a new phylogenetic method for comparing microbial communities. Appl Environ Microbiol. 2005; 71(12):8228–35.

Majumder, R., Sutcliffe, B., Taylor, P. W., & Chapman, T. A. (2019). Next-Generation Sequencing reveals relationship between the larval microbiome and food substrate in the polyphagous Queensland fruit fly. Scientific reports, 9(1), 1–12.

Malacrinò A, Campolo O, Medina RF, Palmeri V. Instar-and host-associated differentiation of bacterial communities in the Mediterranean fruit fly Ceratitis capitata. PloS One. 2018; 13(3):e0194131.

Marín-Cevada V, Caballero-Mellado J, Bustillos-Cristales R, Muñoz-Rojas J, Mascarúa-Esparza MA, Castañeda-Lucio M, López-Reyes L, Martínez-Aguilar L, Fuentes-Ramírez LE. Tatumella ptyseos, an unrevealed causative agent of pink disease in pineapple. J Phytopathol. 2010; 158(2):93–9.

Martin M. Cutadapt removes adapter sequences from high-throughput sequencing reads. EMBnet J. 2011; 17(1):10–2.

Martinson, V. G., Douglas, A. E., & Jaenike, J. (2017). Community structure of the gut microbiota in sympatric species of wild Drosophila. Ecology Letters, 20(5), 629–639.

Martinez-Sañudo I, Simonato M, Squartini A, Mori N, Marri L, Mazzon L. Metagenomic analysis reveals changes of the Drosophila suzukii microbiota in the newly colonized regions. Insect sci. 2018; 25(5):833–46.

Matthies C, Springer N, Ludwig W, Schink B. Pelospora glutarica gen. nov., sp. nov., a glutarate-fermenting, strictly anaerobic, spore-forming bacterium. Int J Syst Evol Microbiol. 2000; 50(2):645–8.

McMurdie PJ, Holmes S. phyloseq: An R package for reproducible interactive analysis and graphics of microbiome census data. PLoS One. 2013; 8(4), e61217.

McMurdie PJ, Holmes S. Waste not, want not: why rarefying microbiome data is inadmissible. PLoS Comput Biol. 2014; 10(4):e1003531.

Morotomi M, Nagai F, Watanabe Y, Tanaka R. Succinatimonas hippei gen. nov., sp. nov., isolated from human faeces. Int J Syst Evol Microbiol. 2010; 60(8):1788–93.

Oksanen J. Vegan: an introduction to ordination. URL http://cran.r-project.org/web/packages/vegan/vignettes/introvegan.pdf. 2015; 8:19.

R Core Team. R: A language and environment for statistical computing. R Foundation for Statistical Computing, Vienna, Austria. 2020; URL R-project.org/.

Rassati D, Marini L, Malacrinò A. Acquisition of fungi from the environment modifies ambrosia beetle mycobiome during invasion. PeerJ. 2019; 7:e8103.

Raza MF, Yao Z, Bai S, Cai Z, Zhang H. Tephritidae fruit fly gut microbiome diversity, function and potential for applications. Bull. Entomol. Res. 2020; 110:423–347.

Riley AB, Kim D, Hansen AK. Genome sequence of “Candidatus Carsonella ruddii” strain BC, a nutritional endosymbiont of Bactericera cockerelli. Genome Announc. 2017; 5(17).

Rivas-Marín E, Devos DP. The Paradigms They Are a-Changin’: past, present and future of PVC bacteria research. Antonie Van Leeuwenhoek. 2018; 111(6):785–99.

Rivers AR, Weber KC, Gardner TG et al. ITSxpress: Software to rapidly trim internally transcribed spacer sequences with quality scores for marker gene analysis. F1000Res. 2018; 7:1418 (doi: 10.12688/f1000research.15704.1)

Rizzi A, Crotti E, Borruso L, Jucker C, Lupi D, Colombo M, Daffonchio D. Characterization of the bacterial community associated with larvae and adults of Anoplophora chinensis collected in Italy by culture and culture-independent methods. BioMed Res. Int. 2013; 2013.

Rota-Stabelli O, Blaxter M, Anfora G. Drosophila suzukii. Current Biology. 2013; 23(1):R8–9.

Silva-Soares NF, Nogueira-Alves A, Beldade P, Mirth CK. Adaptation to new nutritional environments: larval performance, foraging decisions, and adult oviposition choices in Drosophila suzukii. BMC Ecol. 2017; 17(1):1–3.

Solomon, G. M., Dodangoda, H., McCarthy-Walker, T., Ntim-Gyakari, R., & Newell, P. D. (2019). The microbiota of Drosophila suzukii influences the larval development of Drosophila melanogaster. PeerJ, 7, e8097.

Stackebrandt ER, Hespell R. The family succinivibrionaceae. The prokaryotes. Springer, Berlin, Heidelberg. 2006; 3:419–29.

Starmer WT, Fogleman JC. Coadaptation of Drosophila and yeasts in their natural habitat. J Chem Ecol. 1986;12(5):1037–55.

Sudarshan A. Shetty, & Leo Lahti. (2020, October). microbiomeutilities: Utilities for Microbiome Analytics.

Thao ML, Moran NA, Abbot P, Brennan EB, Burckhardt DH, Baumann P. Cospeciation of psyllids and their primary prokaryotic endosymbionts. Appl Environ Microbiol. 2000; 66(7):2898–905.

Tissot M, Stocker RF. Metamorphosis in Drosophila and other insects: the fate of neurons throughout the stages. Prog. Neurobiol. 2000; 62(1):89–111.

Vacchini, V., Gonella, E., Crotti, E., Prosdocimi, E.M., Mazzetto, F., Chouaia, B., Callegari, M., Mapelli, F., Mandrioli, M., Alma, A. and Daffonchio, D., (2017). Bacterial diversity shift determined by different diets in the gut of the spotted wing fly Drosophila suzukii is primarily reflected on acetic acid bacteria. Environmental microbiology reports, 9(2), 91–103.

van Veelen HP, Salles JF, Matson KD, van der Velde M, Tieleman BI. Microbial environment shapes immune function and cloacal microbiota dynamics in zebra finches Taeniopygia guttata. Animal Microbiome. 2020; 2:1–7.

Walsh DB, Bolda MP, Goodhue RE, Dreves AJ, Lee J, Bruck DJ, Walton VM, O’Neal SD, Zalom FG. Drosophila suzukii (Diptera: Drosophilidae): invasive pest of ripening soft fruit expanding its geographic range and damage potential. J Integr Pest Manag. 2011; 2(1):G1–7.

Wang, Q, G. M. Garrity, J. M. Tiedje, and J. R. Cole. 2007. Naïve Bayesian Classifier for Rapid Assignment of rRNA Sequences into the New Bacterial Taxonomy. Appl Environ Microbiol. 73(16):5261–7.

Whitehead SR, Bowers MD. Chemical ecology of fruit defence: synergistic and antagonistic interactions among amides from Piper. Funct Ecol. 2014; 28(5):1094–106.

Yao, Z., Ma, Q., Cai, Z., Raza, M.F., Bai, S., Wang, Y., Zhang, P., Ma, H. and Zhang, H., (2019). Similar shift patterns in gut bacterial and fungal communities across the life stages of Bactrocera minax larvae from two field populations. Frontiers in Microbiology, 10, 2262.

