## Supplementary file for "More persistent bacterial than fungal associations in the microbiota of a pest"

- Supplementary figures

Detailed composition of the fly food medium

| | | amounts per liter | |
|---|---|---|---|
| Water | - | 1000 | mL |
| agar | - | 10 | g |
| sucrose | - | 15 | g |
| glucose | - | 30 | g |
| cornmeal | - | 15 | g |
| wheat germ | - | 10 | g |
| soy flour | - | 10 | g |
| molasses | - | 30 | g |
| Yeast | - | 35 | g |
| Propionic acid | - | 5 | mL |
| tegosept | - | 2 | g |
| ethanol | - | 10 | mL |

- Primers targeting V4V6 region.

Forward primer:
GTGCCAGCMGCCGCGGTAA
Reverse primer:
CGACRRCCATGCANCACCT

- Primers targeting ITS1 region.

Forward primer:
CTTGGTCATTTAGAGGAAG*TAA
Reverse primer:
GCTGCGTTCTTCATCGA* TGC

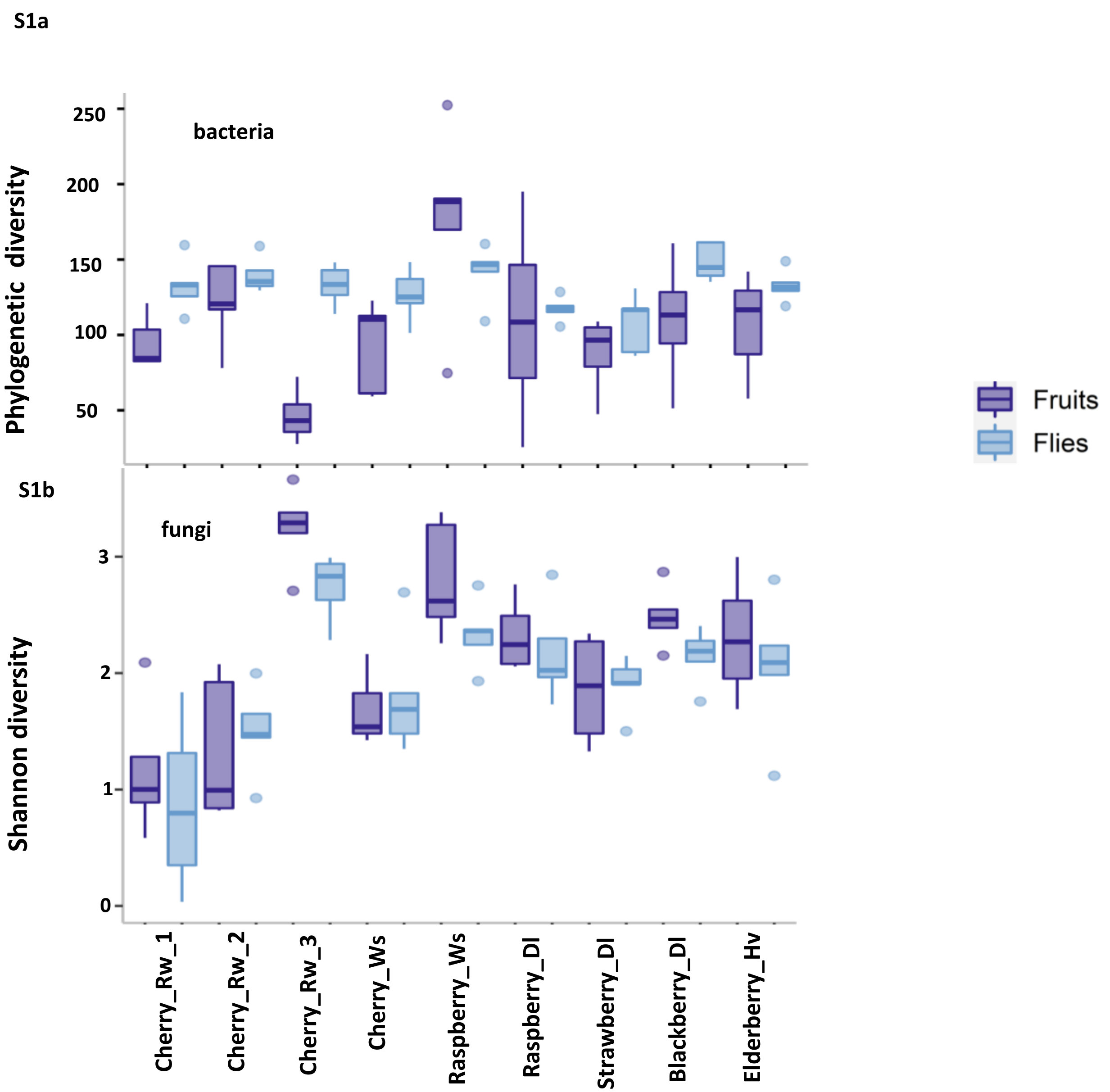


Figure S1a & S1b: Phylogenetic diversity of bacterial communities in fruit and wild fly samples (p-value<0.05) & Shannon diversity of fungal communities in fruit and wild fly samples (p-value <0.001)

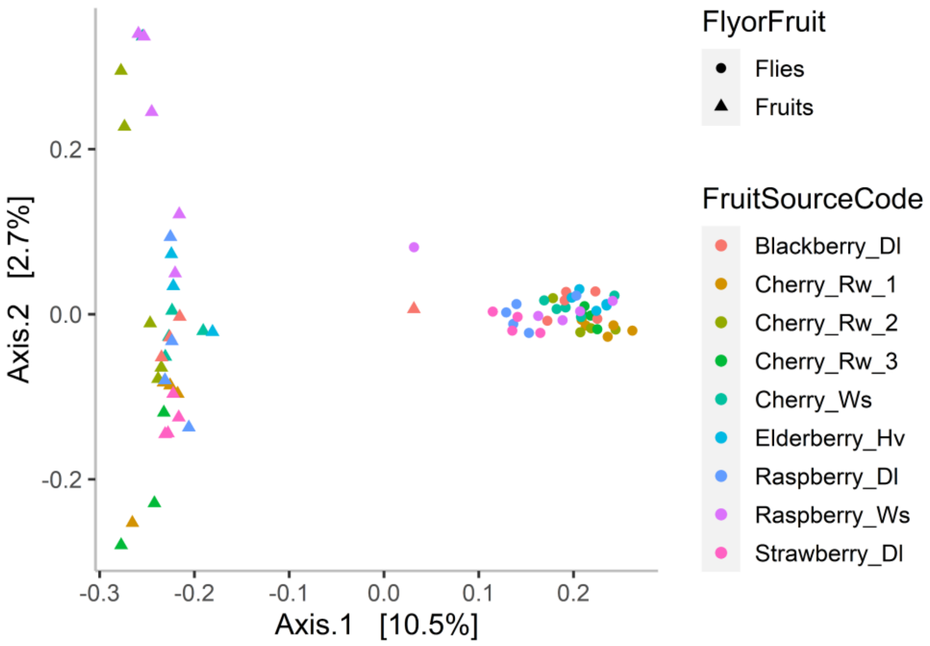


Figure S2: Unweighted UniFrac metrics of bacterial communities in fruits and wild flies (p-values=0.001)

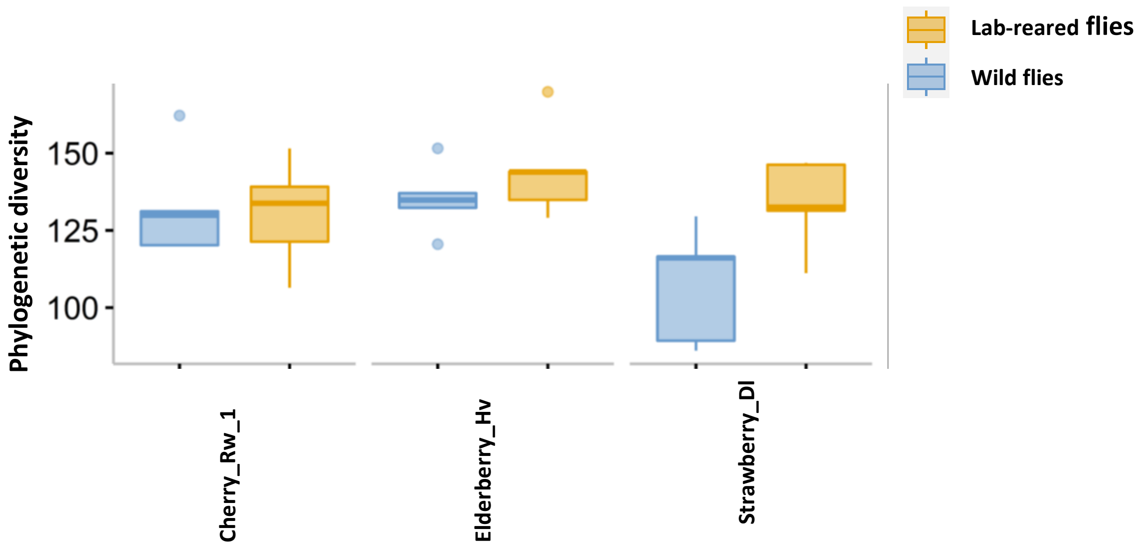


Figure S3: Faith's phylogenetic diversity of bacterial communities in lab-reared and wild flies (p-values=0.1)

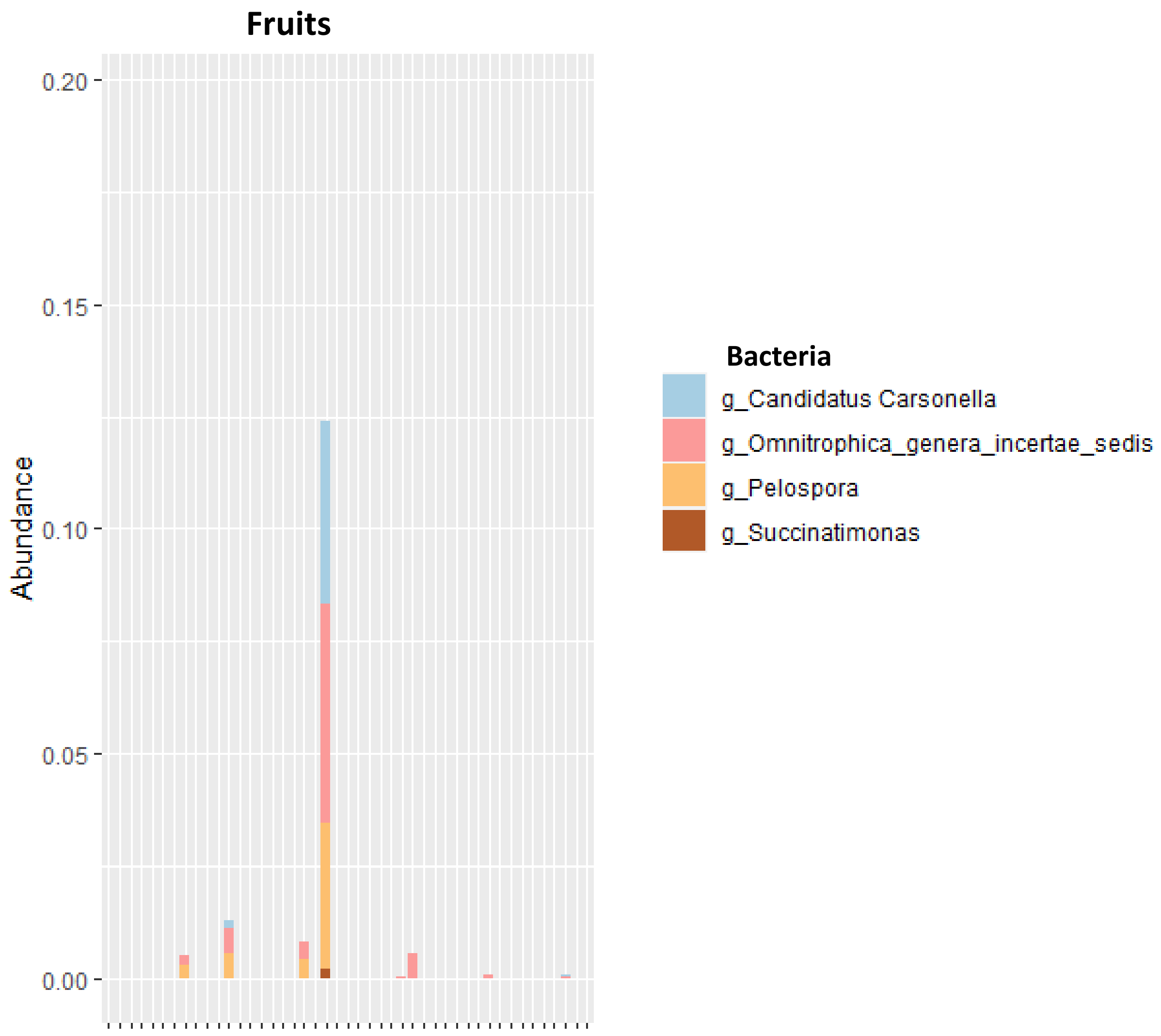


Figure S4: Core bacteria as observed in the fruit samples

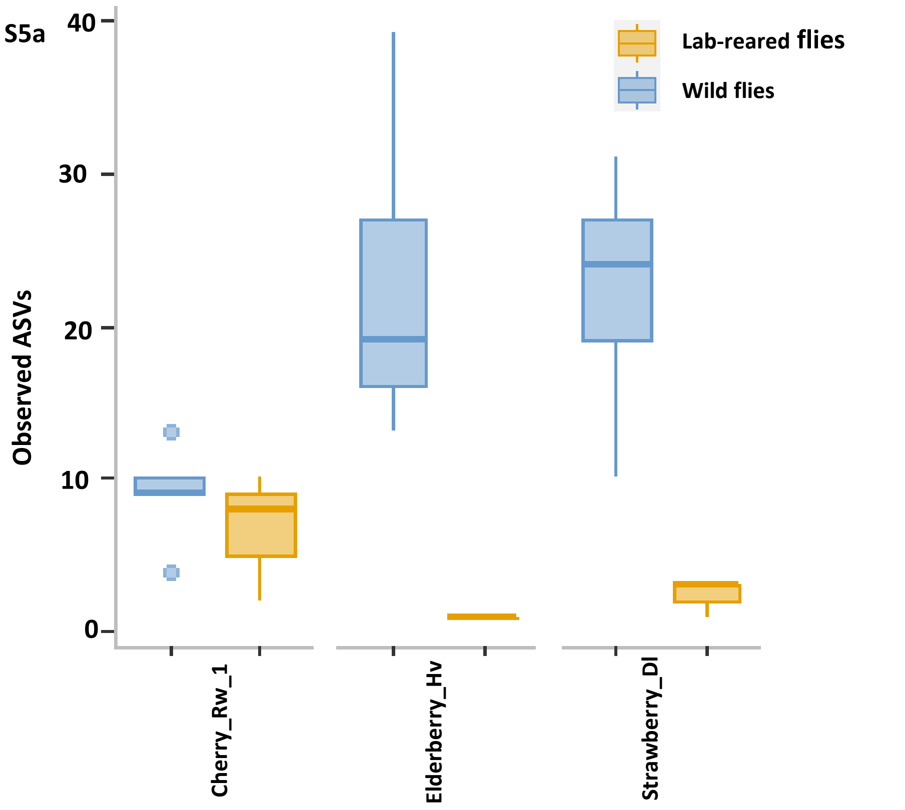


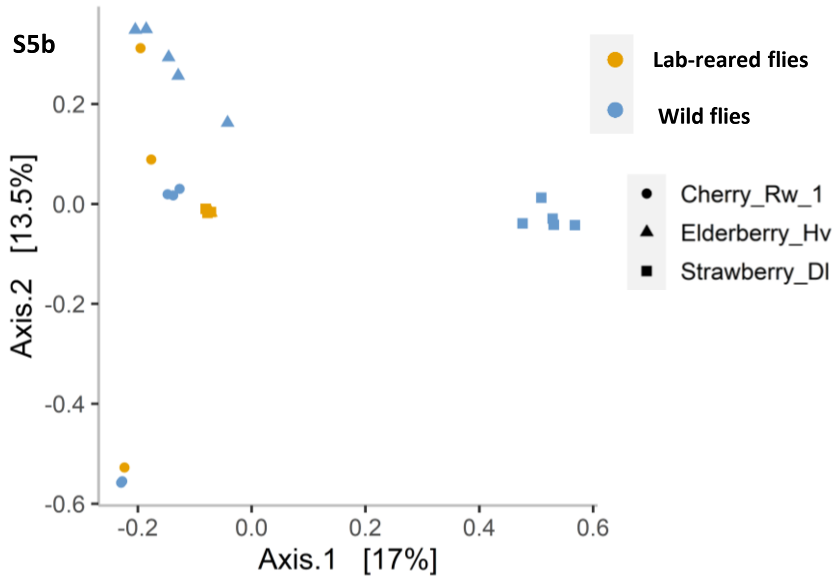


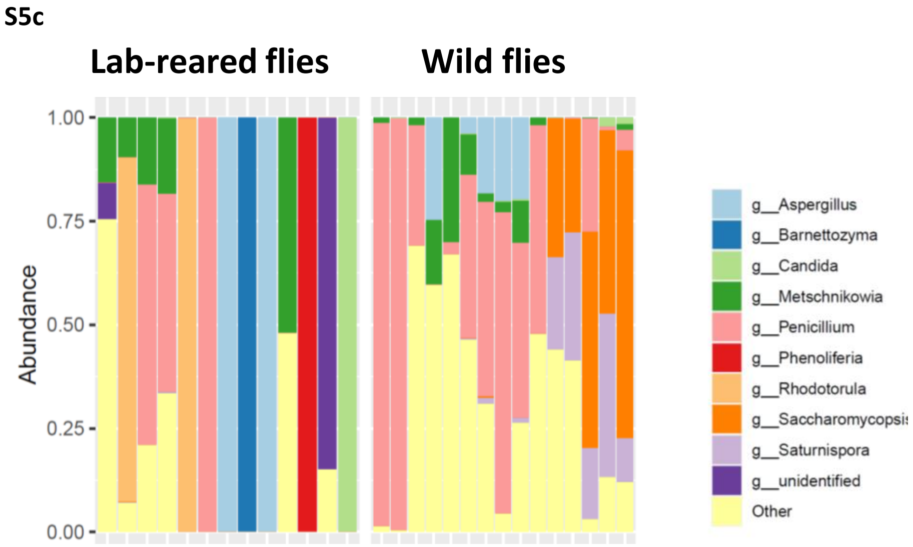


Figure S5: Fungal communities in lab-reared as well as the wild flies. Observed ASVs in lab-reared and wild flies ($X^2$=11.19, p-value<0.001; S5a). Bray Curtis plot shows beta diversity of the fungal communities across lab-reared and wild flies (PERMANOVA, p-value=0.06; S5b). Top 10 abundant fungi as noted across the lab-reared and wild flies (S5c).
